# Dissociable components of oscillatory activity underly information encoding in human perception

**DOI:** 10.1101/2020.09.10.291294

**Authors:** Diego Vidaurre, Radoslaw M. Cichy, Mark W. Woolrich

**Affiliations:** Center for Functionally Integrative Neuroscience, Aarhus University, Denmark, 8000; Department of Psychiatry, University of Oxford, UK, OX37JX; Wellcome Trust Center for Integrative Neuroimaging, University of Oxford, UK, OX37JX; Department of Education and Psychology, Freie Universität Berlin, Germany, 14195

## Abstract

Brain decoding can predict visual perception from non-invasive electrophysiological data by combining information across multiple channels. However, decoding methods typically confound together the multi-faceted and distributed neural processes underlying perception, so it is unclear what specific aspects of the neural computations involved in perception are reflected in this type of macroscale data. Using MEG data recorded while participants viewed a large number of naturalistic images, we analytically separated the brain signal into a slow 1/f drift (<5Hz) and a oscillatory response in the theta frequency band. Combined with a method for capturing between-trial variability in the way stimuli are processed, this analysis revealed that there are at least three dissociable components that contain distinct stimulus-specific information: a 1/f component, reflecting the temporally stable aspect of the stimulus representation; a global phase shift of the theta oscillation, related to differences in the speed of processing between the stimuli; and differential patterns of theta phase across channels, likely related to stimulus-specific computations. We demonstrate that common cognitive interpretations of decoding analysis can be flawed if the multicomponent nature of the signal is ignored, and suggest that, by acknowledging this fact, we can provide a more accurate interpretation of commonly observed phenomena in the study of perception.

## Introduction

In recent years, multivariate pattern analysis (MVPA) or decoding analysis has been successfully used in different cognitive domains to interrogate when and where in the brain information is processed (Norman et al., 2006; Haynes and Rees, 2006; Tong et al., 2012; Haxby et al., 2014; Grootswagers et al., 2017; Cichy et al., 2014; Kragel et al., 2018; Vidaurre et al., 2019). These approaches are typically designed with the objective of maximising the prediction of information, so that it is not always transparent what elements of the data are responsible for the significant predictions. In particular, electrophysiological data have diverse elements, including ongoing oscillations (with amplitude and phase modulations in different frequencies; Buzsaki and Draguhn, 2004), ultra-slow 1/f drifts of activity and slow cortical potentials (He, 2014), non-oscillatory transient changes in signal amplitude (Jones, 2016), cross-frequency coupling (Jensen and Colgin, 2007), and high-frequency burst events (Buzsaki and Lopez da Silva, 2004). Data-driven decoding methods can incorporate all or some of these elements in their prediction, but it is unclear which ones are relevant and how. Because these elements are thought to underpin distinct brain mechanisms, knowing which of them are important for prediction would directly provide important theoretical information about human brain function.

Investigating MEG data recorded while participants viewed object images, we separated task evoked responses into an ultra-slow 1/f drift (<5Hz) and an oscillatory component in the theta frequency band. We then demonstrated their distinct contribution to the multivariate decoding accuracy of image prediction. In particular, isolating the oscillatory component allowed us to extract stimulus-locked phase information, revealing that the phase of the oscillatory component contains critical information about which image category is being processed by the brain. While event-related potentials/fields (ERP/F; Pfurtscheller and Lopes da Silva, 1999) represent stimulus-locked phase information on a single channel basis, here we show that the phase information necessary to decode the stimulus is multivariate across channels. Critically, we show that stimulus-locked phase information can be predictive of the stimulus in two different ways: through differences in the latency of the response that are common across channels; and through relative phase differences between channels, to which single-channel analyses like ERP/F are blind. We argue that these two (coexisting) phase-related features potentially speak to different physiological mechanisms.

In summary, we show that the dissociation of these components provides a more accurate interpretation of commonly observed phenomena in the analysis of perception, in particular with regards to how the stimulus representations generalise (or change) across time points. Further, we provide a better understanding on how multivariate phase information can encode different visual contents in electrophysiological data, and demonstrates how MVPA can be usefully extended to provide interpretable insights on how the brain encodes and processes mental representations.

## Results

We used one of the magnetoencephalography (MEG) data sets presented in Cichy, Pantazis and Oliva (2016), where 118 different image categories were presented 30 times each to 15 subjects. Presentation lasted 500ms, and trials were ~1s long. The multi-channel sensor-space data, epoched around the presentation of each visual stimulus, can be used to train a decoder to predict which visual image is being presented (Cichy et al, 2016).

We reasoned that different aspects of decoding accuracy may separately emerge from different components of the evoked responses. Therefore, we analytically separated the data into two uncorrelated components: an oscillatory component with a dominant theta frequency (6-8Hz); and a slow 1/f component that cannot be considered oscillatory within the trial duration (although it could be related to very slow oscillations in the context of the entire recording); see Methods for details on the preprocessing and organisation of the data, as well as on the decoding methods. **Fig 1A** shows the original signal (low-pass filtered to 10Hz) and the two components for a single trial and channel. **Fig 1B** shows the spectral profile of these two components, averaged across channels and participants. The oscillatory component has a dominant theta frequency, at 6-8Hz. As mentioned, the 1/f component is very slow (in the delta frequency or slower), and not oscillatory at least when considered in 1s trial lengths, as it is suggested by the fact that the spectral profile has no peak.

**Fig 1.**
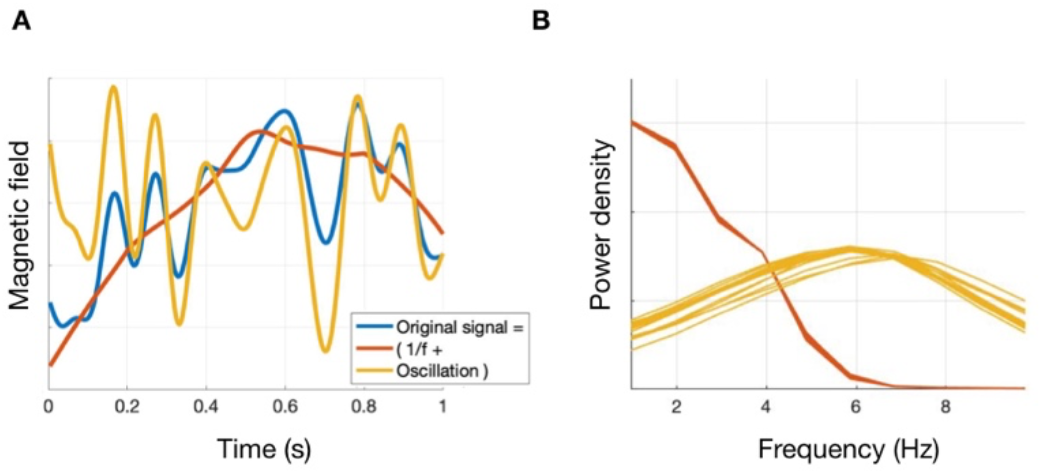
Analytical separation of the signal into oscillatory and a 1/f component. **A**: One trial example. **B**: Power of the original signal and the two components.

### Oscillations can theoretically contain different types of stimulus-specific information

When looking at the oscillatory component, both phase and amplitude modulations can cause differences that can in principle be used by MVPA. Focusing first on phase, we demonstrate schematically different ways by which an oscillatory component might contain stimulus-specific information, either using single MEG channels at a time, or multivariately using multiple channels. This forms the theoretical basis for the subsequent analysis of the importance of phase for MVPA in empirical data.

**Fig 2A** illustrates three ways in which differences in evoked responses could cause changes in decoding accuracy between experimental conditions (i.e., here object images) at the single-channel level. The evoked responses (i.e. the ERP/F) for a single channel are depicted for two different stimuli in red and blue. These differences, examined at two time points that are marked as vertical lines (separated according to the period of the oscillation), and interpreted as differences in decoding regression coefficients (*β*), can either be a phase modulation or an amplitude modulation. When the difference is a phase modulation (top panel), the two stimuli can be distinguished as the blue stimulus evokes an earlier response. As shown in the panels, *β* would have opposite signs between the two time points, which, as we will discuss, is relevant in interpreting decoding accuracy. However, when the difference is an amplitude modulation, two different possibilities exist. The first possibility (middle panel) is a decrease in power, such that *β* would again have opposite signs between the two marked time points. The second possibility (bottom panel) is an additive shift in the signal, which does not cause a sign inversion between the two time points. Therefore, when considered univariately, both phase and amplitude can bring about decoding accuracy.

**Fig 2.**
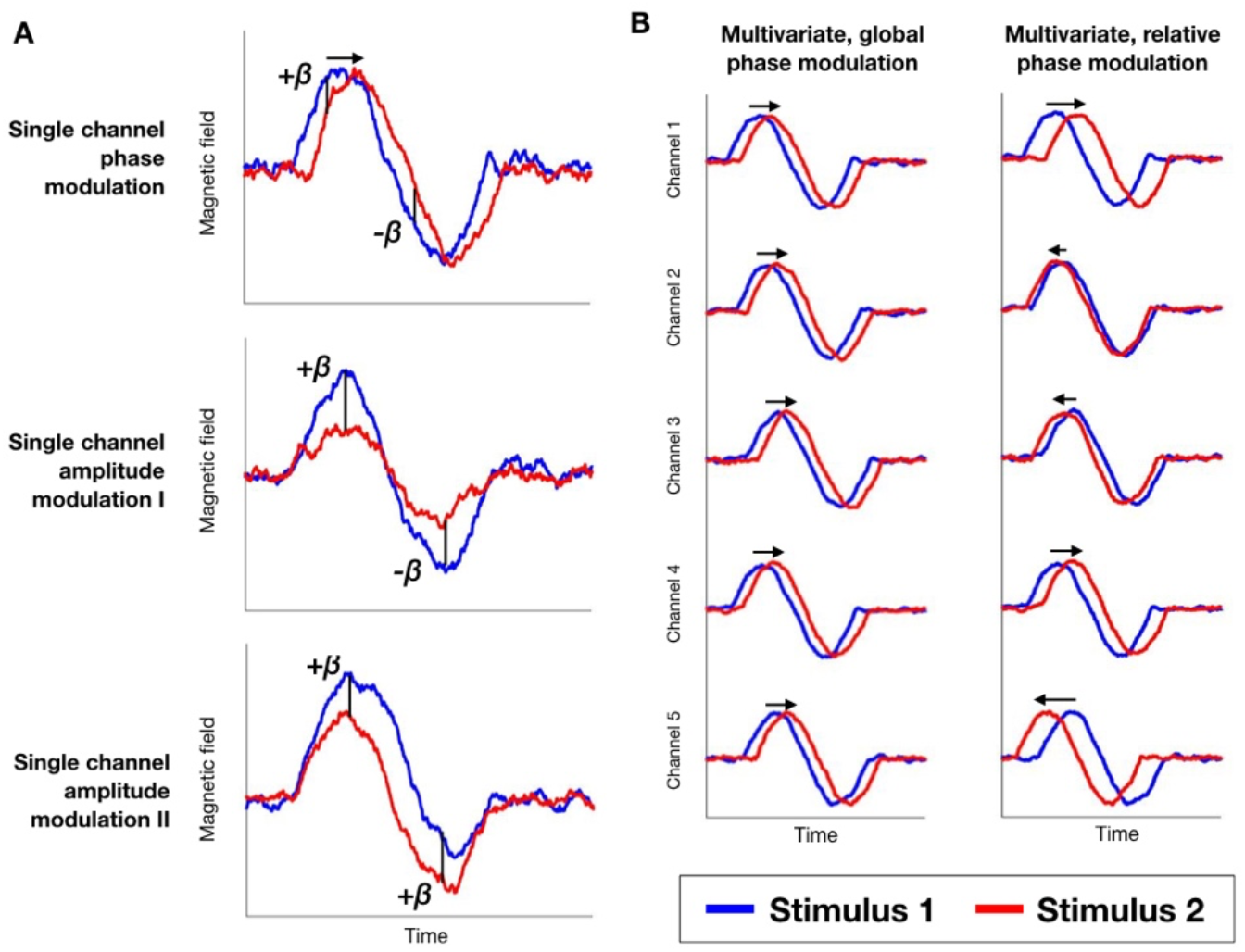
Different types of modulation can bring about decoding accuracy. This can occur either in phase or in amplitude; and either univariately (**A**) or multivariately (**B**). Focusing on phase at the multivariate level, these differences can be either global (consistent across channels) or local/relative (different, between channels). Arrows represent phase differences in phase-specific panels.

Assuming for simplicity that all channels share the same kind of evoked response, **Fig 2B** illustrates two ways in which differences in evoked responses could cause changes in decoding accuracy at the multi-channel level. The horizontal arrows reflect the phase difference between the two stimuli. In one type, the latency of the response is different between the stimuli, such that the phase modulation occurs earlier for some stimuli than for others, but, critically, this delay is global, i.e. consistent across channels (**Fig 2B**, left panel – where all arrows have the same length and orientation). In the other type, it is the local, relative differences in phase between channels that discriminates between stimuli (**Fig 2B**, right panel – where arrows have different lengths and orientations), without there necessarily being a global temporal shift that is common to all channels; i.e. the arrows could sum up to zero across channels.

Importantly, these two different multivariate patterns may afford very different physiological interpretations. The first type of multivariate phase modulations would speak to systematic differences in the timing of information processing; for example, if images of animate beings take less time to process in the brain, then the cascade of stimulus processing would progress faster and that would bring about a change in the time course of the decoding accuracy. The second type of multivariate phase modulations would instead refer to differential phasic patterns between the image categories above and beyond their speed of processing, which could be related to detectable differences in the neural coding of the images.

### Decoding accuracy differentially emerges from dissociable 1/f and oscillatory components

We next show empirically that the oscillatory and the 1/f components contain complementary information underlying discriminability in MVPA of electrophysiological data. Furthermore, we show how some of the most prominent phenomena commonly observed in decoding analysis can parsimoniously be explained by accounting for the separate contributions of these two components. We undertake this in the context of the temporal generalisation matrix (TGM; see Methods), which extends conventional temporally-defined decoding to describe, not just the decodability of the stimulus time point by time point, but also how these neural representations change (or persist) across time points (King and Dehaene, 2014).

**Fig 3A** shows for one representative subject the TGM for the original signal, the 1/f component, the oscillatory component, and the power envelope of the oscillatory component (i.e. with no phase information). For reference, **Fig SI-1** shows the time-resolved decoding accuracy (i.e. the diagonal of the TGM) for one subject and averaged across subjects. The original signal’s decoding accuracy reflects a mixture from the decoding accuracy of the 1/f and the oscillatory components: a linear regression model (done separately for each subject) can predict the original signal’s TGM from the 1/f and the oscillatory components’ TGMs with 67% of explained variance on average across subjects. When using only the 1/f, the explained variance was 9%; and when using only the oscillatory component, it was 59%. This suggests that both components are relevant for decoding in a complementary manner. Importantly, a model including the 1/f component’s, the oscillatory component’s and the power’s TGMs explains on average 7% more variance of the original signal’s TGM than a model that only includes 1/f and power (permutation testing returns p<0.001 for all subject tests), which is a percentage decrement of 12%. This result highlights the importance of phase in decoding analysis. Expanding on this, in **Supplementary Results** we provide empirical evidence on the stimulus-specificity of phase-locking.

**Fig 3.**
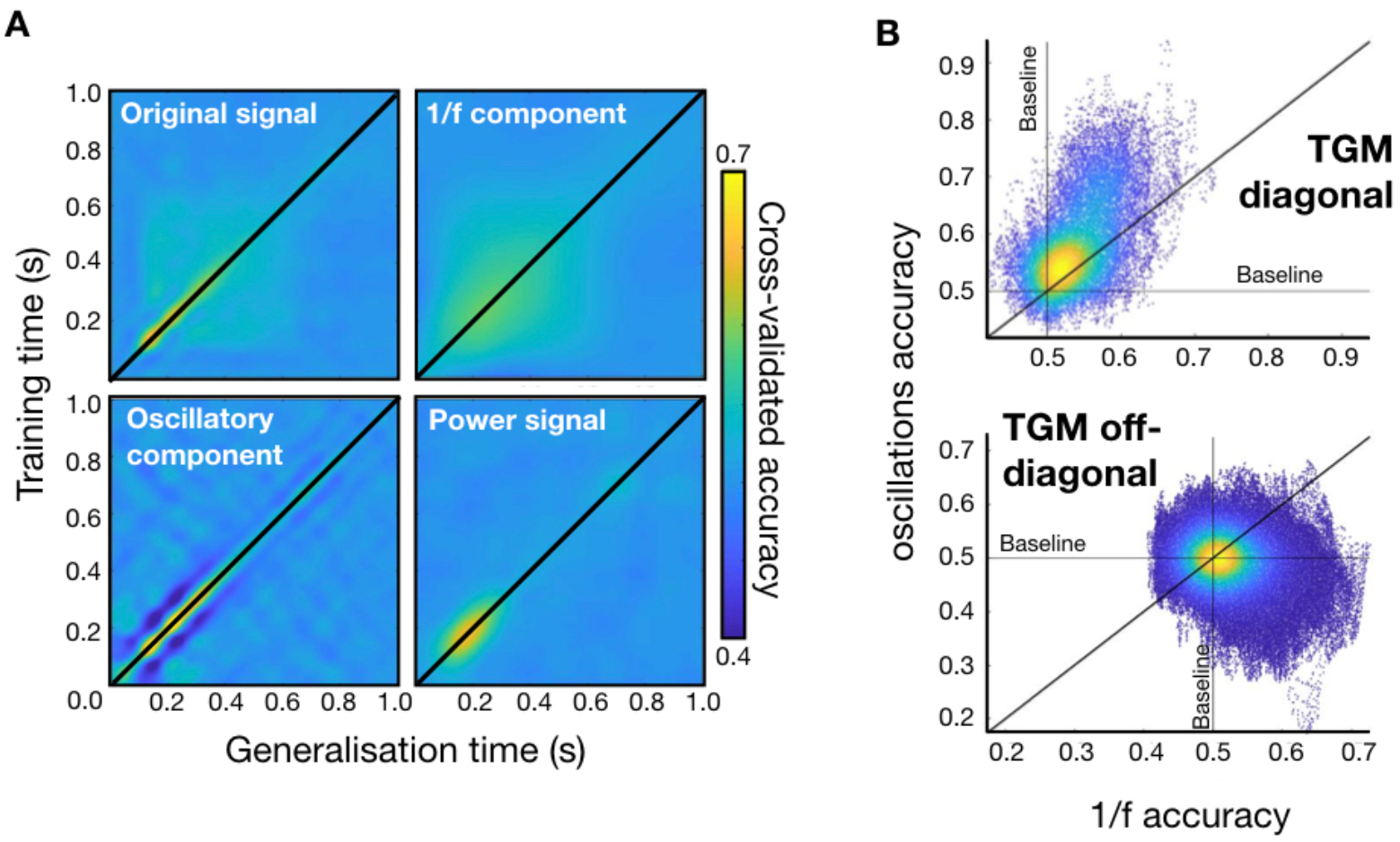
Both 1/f fluctuations and oscillatory activity contribute to MVPA. **A**: TGMs for the original signal, for the 1/f component, for the oscillatory component, and for the power of the oscillatory component. **B**: TGM’s diagonal and TGM’s off-diagonal decoding accuracies of the 1/f component vs oscillatory component. The oscillatory component dominates the diagonal and the 1/f component dominates the off-diagonal of the TGM, indicating that the cross-generalisation decoding accuracy is fundamentally based on the 1/f component, whereas time point by time point decoding accuracy (i.e. the diagonal) is a mixture of both the 1/f and the oscillatory components, but with a larger contribution from the oscillatory component.

**Fig 3B** shows that the contribution to time-resolved decoding accuracy (i.e. the TGM diagonal) is higher for the oscillatory component, which does not generalise well over time; on the contrary, the 1/f component does generalise well over time, contributing more strongly (p-value<0.001; permutation testing) to what is sometimes interpreted as the continuity in time of the neural representation. More specifically, the top panel highlights that the time points of high instantaneous prediction accuracy (i.e. the diagonal of the TGM) are supported by both 1/f and oscillations, but there is a larger contribution of the oscillatory component. The bottom panel suggests that cross-generalising decoding accuracy (i.e. the accuracy that emerges away from the diagonal) is higher for the 1/f component (p-value=0.0182; permutation testing). Thus, this analysis indicates that the 1/f and the oscillatory component contribute to the overall TGM pattern in a complementary way.

Other features of the TGM can be explained by considering the 1/f and the oscillatory component separately. For example, a common pattern observed in decoding analysis is a worse-than-average rebound in decoding accuracy as we get away from the diagonal of the TGM (King and Dehaene, 2014). **Fig 4A** depicts the oscillation-based TGM within the 0.04s to 0.4s interval (left) for one example subject. The right panel represents a section of the matrix at a time point of approximately 200ms and orthogonal to the timeline. Accuracy peaks at the diagonal and suffers a steep decrement below baseline at time points 25-75ms away from the peak. This is consistent with the time occupied by half a cycle of a theta oscillation. We argue that this phenomenon may come about from the oscillatory component with no contribution from the 1/f component. As demonstrated above schematically (top two panels of **Fig 2A**), a rebound in decoding accuracy can come about through either a phase and/or an amplitude modulation causing a sign inversion in the differences in decoding regression coefficients between two time points. Therefore, certain patterns in the oscillatory component, but not in the 1/f, can cause worse than baseline accuracy in the decoding.

**Fig 4.**
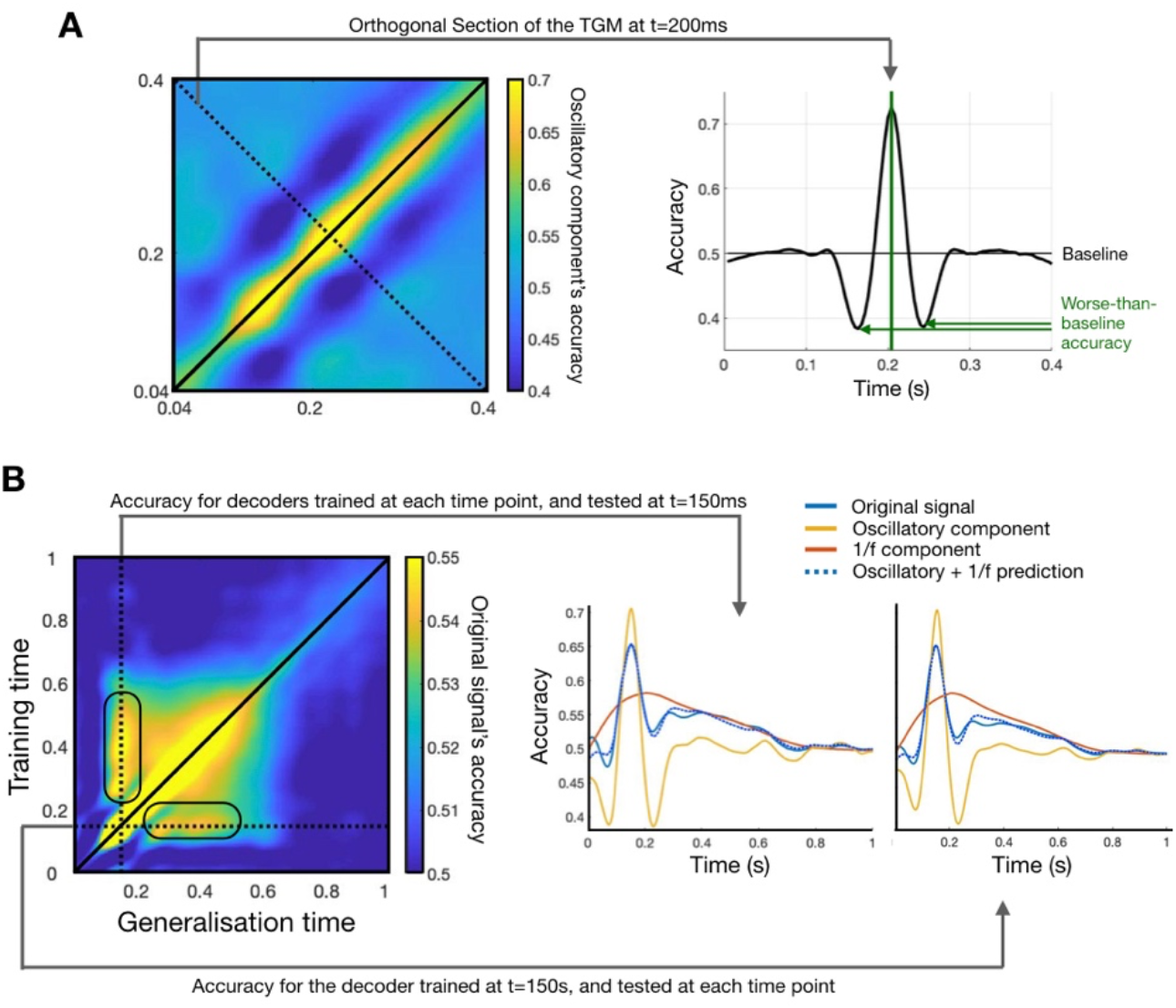
Interpretation of the TGM based on the separation between the oscillatory and 1/f components. **A**: The worse-than-average rebound relates to the oscillatory component. On the left, zoomed-in version of the oscillation-based TGM, within the 40ms to 400ms interval; on the right, a section of the TGM at ~200ms. Result shown for one exemplary subject. **B**. The stronger generalisation at *t*=150ms is due to the combined effect of the 1/f and oscillatory components (see main text). Result shown at the group level.

Another common pattern observed in decoding analysis is a stronger generalisation in regions of the TGM that stem, vertically and horizontally after brief period of depression, from the time points of maximum accuracy in the diagonal – as indicated (at the group level) as rounded boxes in the top left panel of **Fig 4B**. This phenomenon is sometimes interpreted as reactivation of a neuronal representation (King and Dehaene, 2014). Here, we demonstrate that this phenomenon can be explained as an artefact of the manner in which the decoding accuracy provided by the 1/f component and the oscillatory-based phenomenon described in **Fig 4A** combine to bring about the overall decoding accuracy. In particular, we observe that (i) the initial depression of accuracy is caused by the peaks and troughs of the oscillatory component (**Fig 4A**); and that (ii) once such depression is passed, the reactivation is caused by the fact that the 1/f component’s accuracy takes longer to decay than it takes the oscillatory component’s accuracy to come back to baseline. This is shown quantitatively by regressing the 1/f and oscillatory components’ accuracies onto the original signal’s accuracy, the prediction of which is shown as a dotted line in the right panels of **Fig 4B** (explained variance is 96.5% and 97.7%, respectively). Note that the 1/f component’s generalised accuracy has its maximum later than 150ms (i.e. it does better in subsequent time points than at the time point when it was trained); this is only due to the fact that this component’s relevance in the decoding occurs altogether later in the trial. Further note that this reactivation does not manifest in the 1/f and oscillatory components alone (**Fig 3A**), since it is the interaction of both which brings about this effect.

In summary, the key features of the TGM can be parsimoniously explained to a large extent when considering the 1/f and the oscillatory components separately, which should be taken into account when interpreting MVPA results.

### Between-stimuli phase differences are high-dimensional

An assumption often implicitly made when applying MVPA on electrophysiological data is that the relevant information is encoded in multivariate patterns across sensors, rather than in univariate differences, for which single-channel-based analysis would suffice. Here we explicitly test this assumption and ask to what extent there is improved discrimination between the different stimuli when using multivariate (i.e. multi-channel) versus single-channel information (as in ERF/P analysis). That is, given the signal spread in MEG sensor space, would a reduced set of channels be enough to achieve a comparable decoding accuracy?

We know that the signal must be rich in information (either temporally or spatially or both) in order to effectively distinguish between many different stimuli. Also, considering previous work at the level of individual neurons (Stringer et al. 2019) and at the macroscale level in MEG (Nasiotis et al. 2017; Cichy et al., 2015), we hypothesised that the evoked, stimulus-specific activity would be high-dimensional. More specifically, we hypothesised that the stimulus-specific information cannot be encoded in cross-channel averages or other coarse patterns of variation across sensors (such as the first principal component of activity); instead, the information must be encoded in subtle modulations in the multidimensional channel activation patterns.

In order to verify this hypothesis, we conducted a PCA decomposition on the original signal, as well as separately on the 1/f- and oscillation-based components. We kept the first 120 principal components (PC) in each of the three PCA decompositions, which explained around 99% of the variance from the 306 MEG channels. We then computed t-statistics for each PC and pair of stimuli, representing the amount of information each PC carries in isolation about the separation between the two stimuli. **Fig 5** shows, for one exemplary subject, the resulting t-statistics averaged across pairs of stimuli. The fact that the highest t-statistics distribute almost evenly until quite lower PCs (i.e. appearing lower in the matrix and explaining a small amount of variance), highlights the limitations of univariate, ERP/ERF analyses. This is because the information contained in these PCs relates to subtle differences between channels, instead of the main trends of the signal that are shared across channels and could be identified by univariate analyses; such large trends, instead, are contained in the higher PCs.

**Fig 5.**
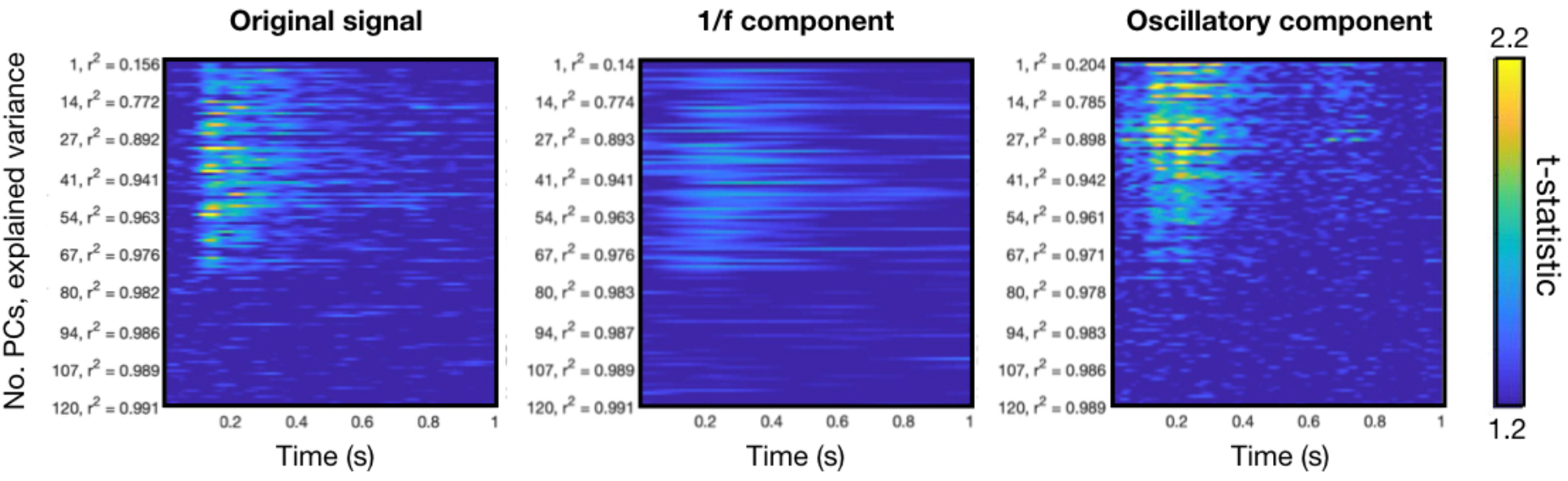
The stimulus-specific information, both for the 1/f and the oscillatory, is high-dimensional. After conducting PCA across channels on the original signal, t-statistics (averaged across pairs of stimuli) express how well each PC distinguish the stimuli for the original signal, the 1/f, and the oscillatory component.

In summary, this analysis shows that the information relevant to stimulus discrimination is high-dimensional, and therefore not solely encoded in the main patterns of covariation of the signal (i.e. the first PCs). This was the case for all aspects of the signal, including the oscillatory component, for which phase is a critical feature. This result has important implications for traditional ERP- and oscillation-based analysis, which are typically based on single channels or averages of channels, highlighting the importance of multivariate analyses.

### Relative vs global phase differences between stimuli: is it just processing speed?

Having demonstrated the importance of the oscillatory component’s high-dimensional phase, we now ask what is the specific nature of the multivariate phase modulation that allows an effective discrimination between stimuli. As discussed above, two non-exclusive possibilities exist. One is that the *global* latency of the response (i.e. common over all sensors) is different between experimental conditions (here stimuli), such that phase-locking occurs a bit earlier for some stimuli than for others and the delay is approximately consistent across channels *(Case 1,* as in **Fig 2B**, left). The other is that the *relative* phase between channels is discriminant between experimental conditions *(Case 2,* as in **Fig 2B**, right), without there necessarily being a global all-channel temporal shift.

To differentiate between these cases, we used the Temporally Unconstrained Decoding Analysis (TUDA) approach presented in Vidaurre et al. (2019) on the oscillatory component of the signal. In brief, TUDA is a Bayesian generative model consisting of a number of decoding models (either classifiers or regressors) together with a data-driven estimation of when each decoding model is active at each trial, i.e. the time courses of each decoding model. The critical feature of TUDA is that the temporal activation of each decoder can be different between trials. Hence, unlike standard decoding, TUDA can effectively accommodate between-trial temporal differences in the stimulus processing cascade. Crucially, these differences are global by definition. By modelling such global differences in the timing of the responses explicitly, TUDA can be used here to separate global from relative differences.

For simplicity and computational efficiency, we trained the model on two supra-level categories into which the stimulus set could be grouped: the size of the stimulus (Size: small, medium, or large), and whether the image corresponds to an animate being (Animacy: yes or no). This way, we could estimate the parameters of the model using all the data for the 118 images and avoided the computational burden of inferring the model parameters for image pair. We set TUDA to use 8 different decoders. **Fig 6A** shows the progression over the time within each trial of the decoders for Size and Animacy, where, for clarity, the trials were ordered according to the first principal component of the decoders activation time courses. To verify the ability of TUDA and standard decoding to fit the data, we conducted permutation testing to assess when TUDA and standard decoding reached a significant prediction, by simply permuting the label of the stimulus and rerunning the models. As shown in **Fig 6B** for one subject (see **Fig SI-3** for all subjects), both TUDA and standard decoding exhibited significant accuracy, even though TUDA had only 8 decoders as opposed to the 250 for standard decoding (i.e. one per time point). This means that we can capture the temporal dynamics of stimulus processing with a relatively low number of parameters, as far as we account for the temporal differences in the data (Vidaurre et al., 2019).

**Fig 6.**
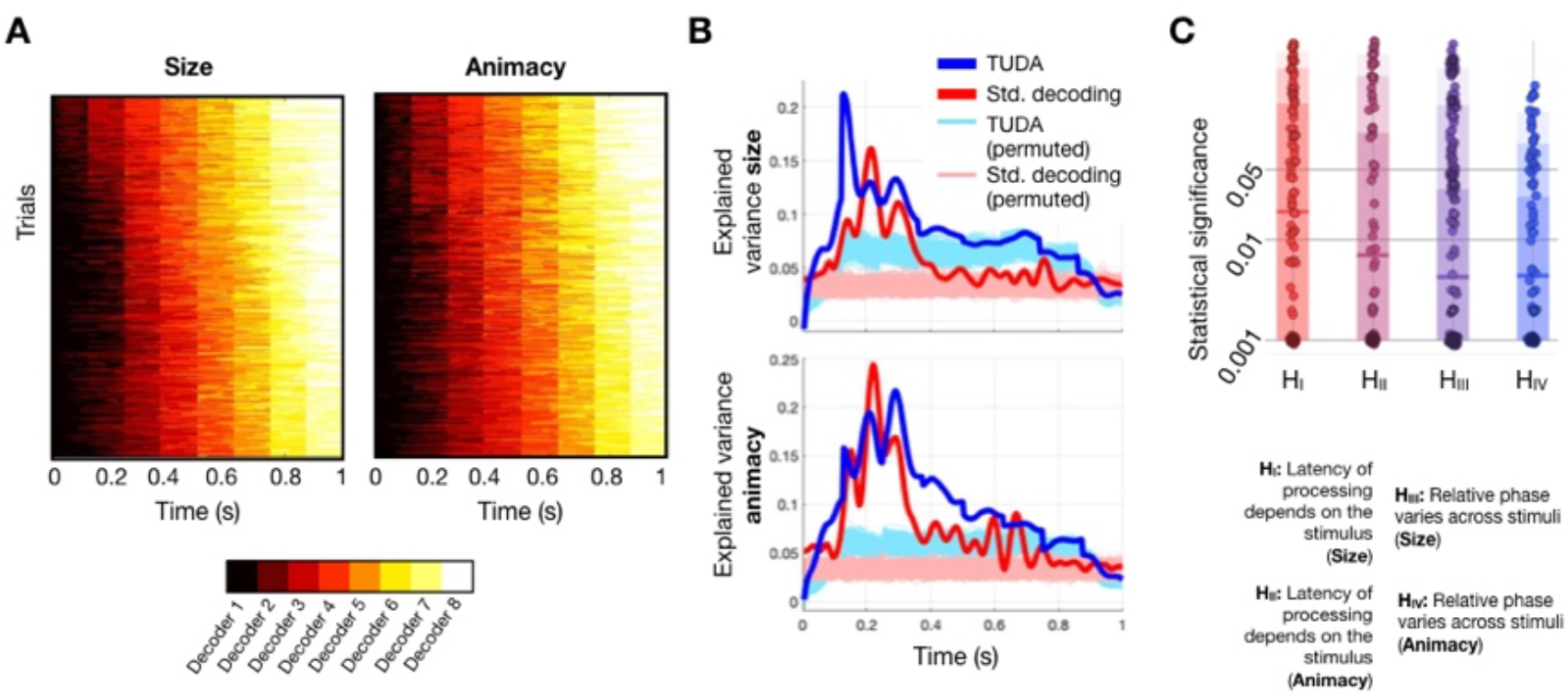
Phasic differences between stimuli in the oscillatory component can be due to global differences in how fast the information is processed for each category, and relative local, phasic differences between channels. **A**: Temporally unconstrained decoding analysis (TUDA) reveals between-trial temporal differences in stimulus processing; the matrices reflect the temporal occupancy of each decoding model for the two considered decoding problems. **B**: Both TUDA and standard decoding exhibit significant decoding accuracy in discriminating animacy and size; statistical significance is given when the accuracy curve is higher than the permutations, represented in lighter colours. **C**: Manhattan plot showing p-values for the hypotheses of whether absolute latency is different between stimuli (size, H_I_; animacy, H_III_), and whether relative latency is different between image categories (size, H_II_; animacy, H_IV_); each dot corresponds to one statistical test for one subject, decoder and pair of category values (e.g. small vs large); the horizontal bars represent the mean of the p-values, and the coloured boxes represents the area containing 90%, 75% and 50% (light, dark, darker, respectively) of the p-values.

By construction, any global (common to all sensors) delays in phase-locking that discriminate between stimuli (Case 1 – **Fig 2B**, left) will be captured in the TUDA model by the trial-specific (and therefore image-category specific) time courses of each decoding model (**Fig 6A**). On these grounds, we used permutation testing to see if differences in the decoders time courses are predictive of Size and Animacy (hypotheses H_I_ and H_III_), yielding one p-value per subject, decoder and pair of image categories. As shown in **Fig 6C**, the majority of tests were significant with a larger effect for Animacy – confirming the existence of global phasic differences between the image categories (Case 1); see Methods for details.

Once the global delays are explained out, all that remains are the relative phase differences between image-categories (Case 2 - **Fig 2B**, left). Therefore, the fact that the decoding models are able to discriminate successfully is evidence of the existence of relative phase differences. To confirm the existence of relative differences, we also asked if phase is more consistent within image categories than it is across image categories *during* the time that each decoder is active. For example, for the time points when, e.g., Decoder 2 is active, we ask if phase-locking is significantly higher within category than between category. If the answer is positive, this would confirm the existence of differential phase patterns between the image categories above and beyond their speed of processing. To quantify phase locking, we used the Phase Locking Factor (PLF; Lachaux et al., 1999), which measures, for any given time point, how similar for each channel are the values of the phase across trials within a certain frequency band (see Methods). We then used permutation testing to assess, for each decoder (i.e. for the time points where each decoder is active), whether PLF was higher within than between image categories. Hypotheses H_II_ and H_IV_ correspond respectively to Size and Animacy, with one test per subject, decoder and pair of image categories. **Fig 6C** shows that there is a majority of significant p-values, especially for Animacy. Overall, the distributions of p-values are strongly skewed towards significance.

In summary, these results suggest the existence of significant phase differences between stimuli at two different levels: global shifts in phase, attributed to the stimuli having diverse processing latencies (Vidaurre et al., 2019); and relative phase differences, attributed to the existence of stimulus-specific, multivariate phase configurations. As mentioned earlier, these two cases may afford very different physiological interpretations.

## Discussion

In recent years, the application of MVPA on electrophysiological recordings has made progress in understanding how the brain implements stimulus processing. However, this progress is hampered by our lack of comprehension of which aspects of the signal drive successful decoding. Because electrophysiological signals are complex and receive influence from multiple brain mechanisms, we argue that we will only be able to further advance our understanding of stimulus processing by breaking down the signals into their underlying constituents and analysing their contribution to decoding both separately and in combination. In this paper, we followed this principle by analysing the separate contributions of the oscillatory (5-10Hz) and the 1/f (<5Hz) components of the signal. We found that the patterns of decoding accuracy commonly observed in the literature can be well-explained from the distinct contributions of these two components, showing that the interpretation of these patterns can be misleading if we do not account for the separability of such contributions. This asks for a reconsideration of the way these patterns are read in the field. For example, certain patterns observed in the TGM are often interpreted as reactivation of mental representations, with no consideration of the fact that these patterns might arise from basic properties of the signal. While the cognitive interpretation is not necessarily incorrect, our statement is that more robust interpretations will emerge from embracing the complexity of the signal and breaking down how these fundamental properties relate to cognition.

Here, we paid special attention to the oscillatory component, analysing how the multivariate phase of electrophysiological recordings contain information about the specific stimulus being processed by the brain. Our analyses stressed the importance of multivariate over purely univariate analyses, supporting the claim that the former are better able to account for systemic nature of the nervous system (Varela et al., 2001; Kragel et al., 2018). Critically, we showed that two dissociable phase-related mechanisms can bring about decoding accuracy: differences in the latency of processing (e.g. some images are processed quicker than others), and differential patterns of phase across channels, which can be attributed to the specifics of each neural representation (Cichy et al., 2015). These two types of effects speak to distinct neural phenomena affording very different cognitive interpretations. Furthermore, their relative influence can determine how MVPA can be used in different scientific contexts. For example, when the timing of the response is uncertain as it is in the case of imagery (given the lack of an external trigger), MVPA needs to rely on relative phase differences given that absolute phase differences are difficult to control for. The fact that decoding has been applied in this setting successfully (Xie et al., 2020; Dijkstra et al. 2020) brings additional evidence on the existence of relative phase differences in visual processing as measured by MEG. In cases where the spatial resolution of MEG might be insufficient to capture fine-grained relative differences, the absolute differences could be the only driver of a successful prediction. This could be the case of pain processing, for instance, where a large part of the relevant signal originates from deep regions that are harder to access by non-invasive electrophysiological modalities (Tracey and Mantyh, 2007).

We note that the analysis performed here was by no means intended to be exhaustive in accounting for all the different features contained in the data. For simplicity, we removed all information content above 10Hz, thereby disregarding higher frequencies such as those in the gamma band, which has purportedly a critical role in cognition (Herrmann et al, 2004). Nor did we look at the how the interactions between the 1/f component and the oscillations is predictive of the perceived stimulus. Future investigations will be needed to understand the role of these other components. We note also that, as opposed to **Fig. 2**, in real data the channels will likely not share the same kind of evoked response, and some channels will show no relevant response at all. This however results in no loss of generality for the above arguments.

An important open question about the mechanistic underpinnings of stimulus processing is whether the stimulus-specific evoked activity is caused by a reorganisation of the ongoing phasic trajectories (i.e. through phase resetting) or is otherwise generated independently of the ongoing phase. Whereas we showed here that there is information contained in the patterns of phase-locking across trials, this does not necessarily imply the existence of phase-resetting (Shah et al. 2004; Mazaheri and Jensen, 2006; Mäkinen et al., 2005). Distinguishing between these two cases is not obvious in non-invasive electrophysiological data (Sauseng et al., 2007), and, at the bare minimum, requires a sufficient baseline period that we lack here. We however speculate that, most likely, the observed effects will not be purely caused by neither phase-resetting nor a separate event-related activation, but instead be some mixture of both, with proportions that depend on the brain region and the type of stimulus (Fell et al., 2004). It is also possible that phase-resetting occurs, but studies focused on univariate analysis could not find it, simply because the effect relies on subtle differences across channels that can only be identified using a multivariate approach.

More broadly, there is the more general question of how the whole-brain ongoing (prestimulus) phase configuration of the brain in slow frequencies influences stimulus processing at higher frequencies. Previously, we showed that there exist reliable whole-brain patterns of phase coherence in theta and alpha at rest (Vidaurre et al., 2018), and that these configurations hold some relation to gamma frequency stimulus responses (Hirschmann et al., 2019). Considering these and other many studies about the influence of the low frequency ranges on diverse aspects of cognition (Baria et al, 2017; Monto et al., 2008; He, 2014; Nobre and van Ede, 2018) and the different ways that ongoing brain states modulate neural responses (Podvalny et al., 2019; McCormick et al., 2020), it is clear that part of the large variability observed in the neural responses could be explained by the ongoing, large-scale network activity and concurrent phase trajectories, and that accounting by these factors could arguably improve the capacity of MVPA to produce successful predictions.

In summary, we propose that macroscale brain signals contain stimulus-specific information at various different levels, and that dissociating them and understanding their potential hierarchical organisation is likely to be the route to eventually breaking down the multifaceted aspects of information processing in the brain.

## Materials and Methods

### Basic theoretical background

Conventional MVPA and event-related potentials/fields (ERPs/ERFs), which represent the average pattern of the signal locked to the presentation of the stimulus (Pfurtscheller and Lopes da Silva, 1999), are both based on assuming consistent timing (or phase) over trials. This is because conventional MVPA uses the same decoder over all trials at the same time-point within each trial, and ERP/ERF approaches average over trials at the same time-point within each trial. The most fundamental differences between these two approaches relies on MVPA being prediction-based (i.e. decoding-based) and, critically, multivariate over channels – while ERP/ERF analysis is univariate over channels and encoding-based (Weichwald et al., 2015).

We examined MVPA in the context of the temporal generalisation matrix (TGM) approach (King and Dehaene, 2014), which extends conventional time-resolved decoding analysis. The TGM, as estimated from magnetoencephalography (MEG) in MVPA approaches, is widely used to assess the dynamics of neural representations, is a T by T matrix of decoding accuracies (where T is the number of time points in the trial), such that one decoding model is trained per time point and tested on each one of the time points of the trial in a cross-validation fashion. The diagonal of the TGM reflects how well can we decode information time point by time point, indicating the emerging and waning of the different stages of stimulus processing. The off-diagonal of the TGM shows how decoding models generalise to different data points of those where they were trained, and, therefore, is argued to reflect the stability of the neural code for the neural representation under study. Thus, the off-diagonal elements of the TGM can be interpreted as related to memory in the most basic definition of the term; that is, the persistency of information and meaning in the brain for longer than an instant. Because it is data-driven, however, the construction of the TGM is not explicitly concerned by the idiosyncrasies of electrophysiological data. The TGM is for example uninformed of how oscillations relate to decoding accuracy. This is in contrast with the extensive body of research on the importance of oscillations in cognition and information representation (Buzsaki and Draguhn, 2004).

### Signal processing

Data were downsampled to 250Hz, and low-pass filtered under 10Hz in order to narrow down the analysis on alpha and lower frequencies. We discarded higher frequencies for simplicity, without any loss of generality in the conclusions.

For each trial and channel, we analytically separated the signal into two uncorrelated components. The first component was computed by applying local regression using weighted linear least squares and a 1^st^ degree polynomial model (parametrised to use 100 points, i.e. 10% of the trial). This yielded a slow component or trend, which we referred to as 1/f. Note that this component is not oscillatory when considered in 1s trials, but it could contain traces of ultraslow oscillations in the context of the entire recording. We then regressed out this component onto the original signal. The residual is a detrended oscillation, which we referred to as the oscillatory component. We used this approach to avoid the sinusoidal assumption of Fourier analysis and the limitations of filtering, which might not be appropriate for the frequencies considered here and the length of the trials (1s).

### Methods for standard decoding analysis

We used cross-validation to assess the accuracy and cross-generalisation accuracy (King and Dehaene, 2014) of the decoding for each pair of stimuli. Given a set of trials or repetitions of the experiment, a decoder (regressor or classifier) contains information about how the stimulus is represented in the brain at the time that the decoder was estimated. Mathematically it can be represented as a function *f*(*x_t_,y_t_*), which predicts the stimulus *y_t_* from the data *x_t_* at time point *t.* As a base decoder we used L2-penalised linear regression, usually referred to as ridge regression (Friedman et al, 2001s), where the predicted variable was encoded as −1 or +1, and we selected the regularization parameter using a nested cross-validation loop. Although logistic regression is more adequate to deal with binary classification, we opted for L2-penalised regression because its lower computational cost, which is relevant here due to the high number of stimuli pairs. Note also that standard linear regression, which is intimately related to L2-penalised linear regression (since they both minimise the sum-of-squared errors), is equivalent to linear discriminant analysis insofar as the proportions of the classes are equal, as occurs here (Friedman et al, 2001).

### Temporally-unconstrained decoding analysis

Let us assume that we can represent how the brain encodes information using a set of *K* decoders, i.e. functions *f_k_* that can predict with certain accuracy the value of the perceived stimuli given the brain data. We will also assume that at any given time point of the experiment, there is only one “active” decoder; or, in other words, that every time point was used to train one and only one decoder *f_k_*. For convenience, let us define an indicator variable *γ_tnk_*, such that *γ_tnk_* = 1 if the *k*-th decoder is active at time point *t* and trial *n,* and *γ_tnk_ =* 0 otherwise.

For standard decoding, the basic premise is that *γ_tnk_* has the same value (either 0 or 1) for all trials *n* = 1 …*N*. Also, typically, there is a different decoder for each time point *t* = 1 …*T*, where *T* is number of time points in each trial. Hence, there are *K* = *T* decoders for standard decoding in total.

In the Temporal Unconstrained Decoding Analysis (TUDA) approach, the key idea is to relax the assumption that *γ_tnk_* has the same value for all trials. In exchange, the number of decoders is considerably lower than with standard decoding (i.e. *K* is much lower than *T*), so that we are trading temporal flexibility for spatial parameters. While the estimations of the decoders in standard decoding are decoupled from each other, TUDA is a unitary Bayesian (generative) model that includes the regression parameters of the *K* decoders, the parameters *γ_tnk_*, and a transition probability matrix *M* containing the probabilities of transitioning from one decoder to another within the trials. Similarly to the standard decoding approach described above, the base decoder used for TUDA is a regularised regression model, which makes the results comparable between the two approaches. All the elements of the TUDA model are estimated using variational Bayesian inference (Vidaurre et al. 2016; Vidaurre et al, 2018b).

Although this model was already presented previously (Vidaurre et al, 2019), here we applied three modification to improve performance and avoid overfitting. Indeed, overfitting can be a problem of this model if some decoders specialise too much in certain stimulus values. For example, in the hypothetical case that there is an overall shift in the signal when face stimuli are presented as opposed to inanimate objects, there would be the risk that a subset of the decoders takes all time points for “face” trials (by having overall higher regressor parameters), while a separate subset of the decoders takes the other trials (by having overall lower regressor parameters). This degenerate solution is of course not useful to interrogate the cascade of stimulus processing. The improvements we have introduced in this paper are:

- The decoders are forced to activate in sequence, so that every trial starts with the first decoder (*k* = 1) and finishes with the last decoder (*k* = *K*).
- The decoders are forbidden to reach out too far in the trial from their mean activation point (*k/T*). In particular, when the decoders are allowed to be active is given by the matrix shown in Fig SI-7.
- To avoid the aforementioned effect, we constrained the estimation of the decoder parameters so that for all decoders the regression parameters sum up to the same value.

Because the inference of this model can take a bit longer than a standard decoder, we applied TUDA on image “supra”-categories: to whether the image is an animate being, and the size of the represented object. This way, we could also use all trials in the estimation.

### Testing the latency of processing and the relative phase between image categories

We used permutation testing for both tests. To test the latency of processing, we took the decoding time courses *γ*, which is a matrix of dimension *T* x *N* x *K* (where *N* is the number of trials), and reorganised it into a matrix *N* x *TK,* such that each row corresponds to one trial, and contains (for that trial) the time course of activation of each decoder. Note that if we did this for standard decoding, all rows would be equal. We then used permutation testing to assess whether we can predict the stimulus using this matrix. If this is the case, then that points at different speeds of processing for each image category.

Now, to test if there are cross-channel differences in the relative phase between image categories, we need to factor out the differences in absolute latency (which were assessed in the previous test). Only then, we will be able to test if the nature of the phase trajectories (i.e. the relative phase between channels at each time point) is distinct across image categories above and beyond how fast these trajectories traverse. We do this by using the temporal realignment provided by the activation times of the decoders, defined by *γ_tnk_*; these were used as data-driven time windows containing quasi-stationary stages of processing, i.e. with a quasistable differential phase pattern. Now, for each of the decoders (i.e. within the decoder-specific time windows), we tested whether the within-class PLF was stronger than the between-classes PLF. We tested this using permutation testing, aggregating the PLF values across channels. If phase locking was found to be more consistent within image categories than in general, that would suggest distinct relative phasic patterns across channels for each stimulus condition.

### Quantifying phase-locking

Given that the distribution of phase across trials in the absence of stimuli is random, we can describe phase-resetting, for each time point after stimulus presentation, as how concentrated is the across-trials distribution of phases at a specific angle. This is commonly measured in practice using the Phase Locking Factor (PLF):

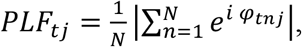

where *φ_tnj_* is the instantaneous phase of trial *n* at time point *t* for channel *j* (Lachaux et al., 1999).

### Code availability

The custom code to reproduce the analysis can be found in the first authors Github personal site. The code for running TUDA can be found in https://github.com/OHBA-analysis/HMM-MAR.

## Supplemental information

### Supplementary Results

#### Phase-locking is stimulus-specific

We have shown that phase of the oscillatory component contains relevant information about the stimuli, and that MVPA can leverage this information to discriminate between the different image classes. We here provide further evidence that this is the case by showing that, not only that there (obviously) must exist at least a certain amount phase-locking for decoding analysis to succeed, but also that the nature of such phase-locking is stimulus-specific and thus is important to decoding.

Phase-locking is necessary because if the phases were completely random across trials, then trials with a negative field would cancel trials with a positive field, with the result that phase-based decoding would be impossible. Phase locking between trials at each time point can be quantified using the Phase Locking Factor (PLF; Lachaux et al., 1999), which measures, for any given time point, how similar for each channel are the values of the phase across trials within a certain frequency band (see Methods). To confirm this, **Fig SI-2A** shows, for one representative subject, the PLF per time point (averaged across channels) together with the time-resolved decoding accuracy (i.e. the diagonal of the TGM); the correlation of these two metrics is 0.94 (0.935 on average across subjects), highlighting the importance of phaselocking for MVPA. Furthermore, we computed a t-statistic 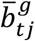 representing how important a given channel *j* is for the discrimination of stimulus *g* at time point *t* (see Methods). More precisely, the “decoding weights” in the figure are univariate t-statistics instead of multivariate regression coefficients, because univariate statistics are more representative of the relevance of a feature than multivariate regression coefficients (Haufe et al 2014). Focusing on the peak of the phase-locking for an exemplary subject (*t*=200ms), **Fig SI-2B** confirms that the channels exhibiting a higher degree of phase coupling are also those that contribute more strongly to the decoding estimation; the scatter plot shows this relationship such that each point corresponds to a particular channel and stimulus. The histogram in **Fig SI-2C** shows the distribution of correlations between the PLF values and the decoding weights across channels, pooled across stimuli and subjects. The correlations are significantly higher than 0.0 (p<0.001, permutation testing) for all subjects.

Having shown that phase-locking is necessary, we further demonstrate that it is also stimulus-specific. Using permutation testing and focusing on *t*=0.2s, we tested whether there is more phase-locking between trials that correspond to the same stimulus than between trials that correspond to different stimuli (see Methods). **Fig SI-2C** presents the p-values across subjects and pairs of stimuli, showing that the timing and nature of such phase-locking is highly stimulus-specific (i.e. most p-values are significant).

### Supplementary Methods

#### Measuring channel relevance for decoding

In order to get an estimation of how important is a channel for the decoding, we can compute, for each channel *j*, time point *t* and pair of stimuli *g* and *h,* an unsigned t-statistic representing how important is that channel for the decoding at *t*:

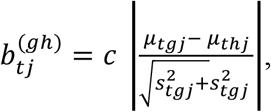

where *μ_tgj_* and 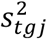 are, respectively, the mean and the variance of channel *j* at time point *t* for stimulus *j*, and *c* is a constant. We then aggregated these for each stimulus, with respect to all the other stimuli, such as:

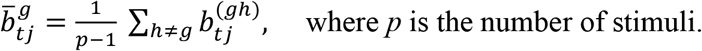

These 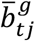 values represent how relevant is channel *j* at time *t* to discriminate stimulus *g*. These were used instead of multivariate decoding weights because of their higher interpretability (Haufe et al, 2014).

## Supplementary Figures

**Fig SI-1.**
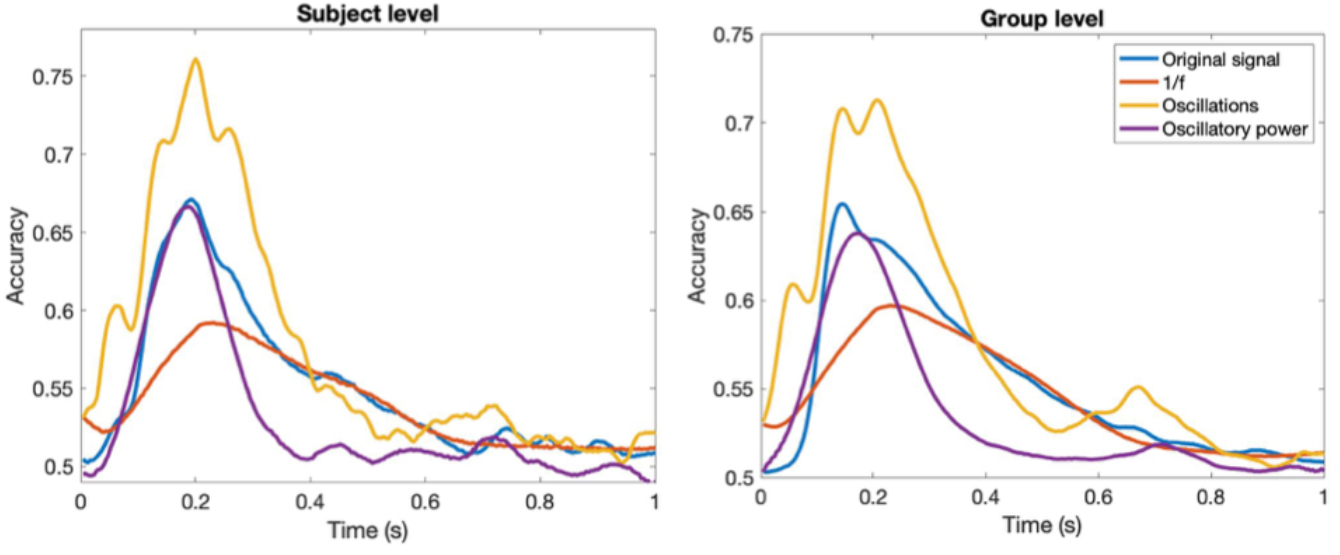
Time-resolved decoding accuracy for one subject and at the group level.

**Fig SI-2.**
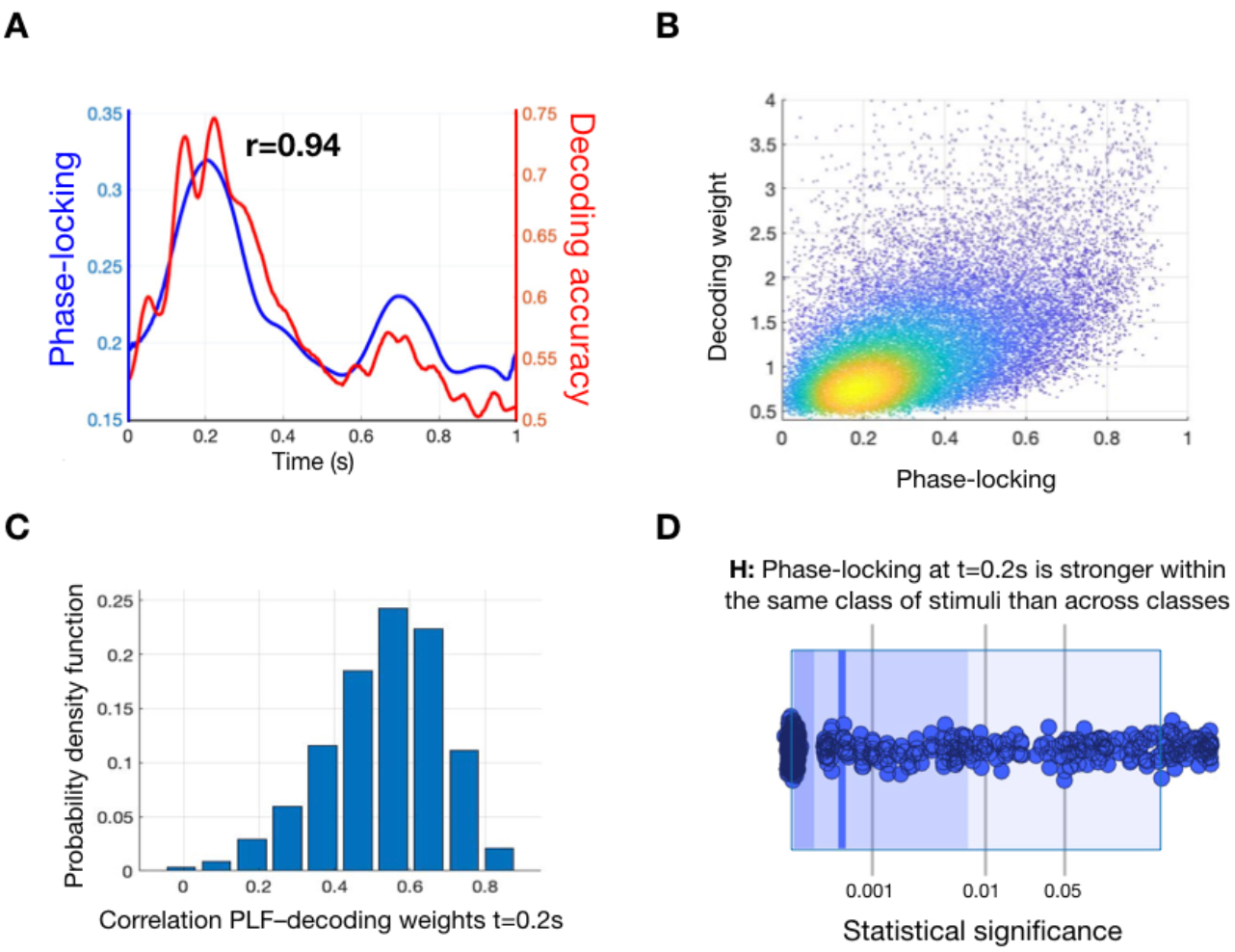
Oscillation-based accuracy necessitates phase-coupling across trials. **A**: The phase-locking factor (PLF; averaged across channels) and oscillation-based decoding accuracy are highly correlated across time points. **B**: Focusing on time point *t*=0.2s, scatter plots of PLF vs decoding weights for one subject. **C**: Histogram of correlations between the two measures across channels, stimuli and subjects (correlations are highly significant, p<0.001). **D**: Distribution of p-values for the hypothesis that phase-locking is stronger within trials belonging to the same stimulus than between different stimulus at *t*=0.2s, indicating that phase-resetting is highly stimulus specific; each dot represents one p-value, the vertical blue bar represents the mean of the p-values, and the coloured boxes represents the area containing 90%, 75% and 50% (light, dark, darker, respectively) of the p-values.

**Fig SI-3.**
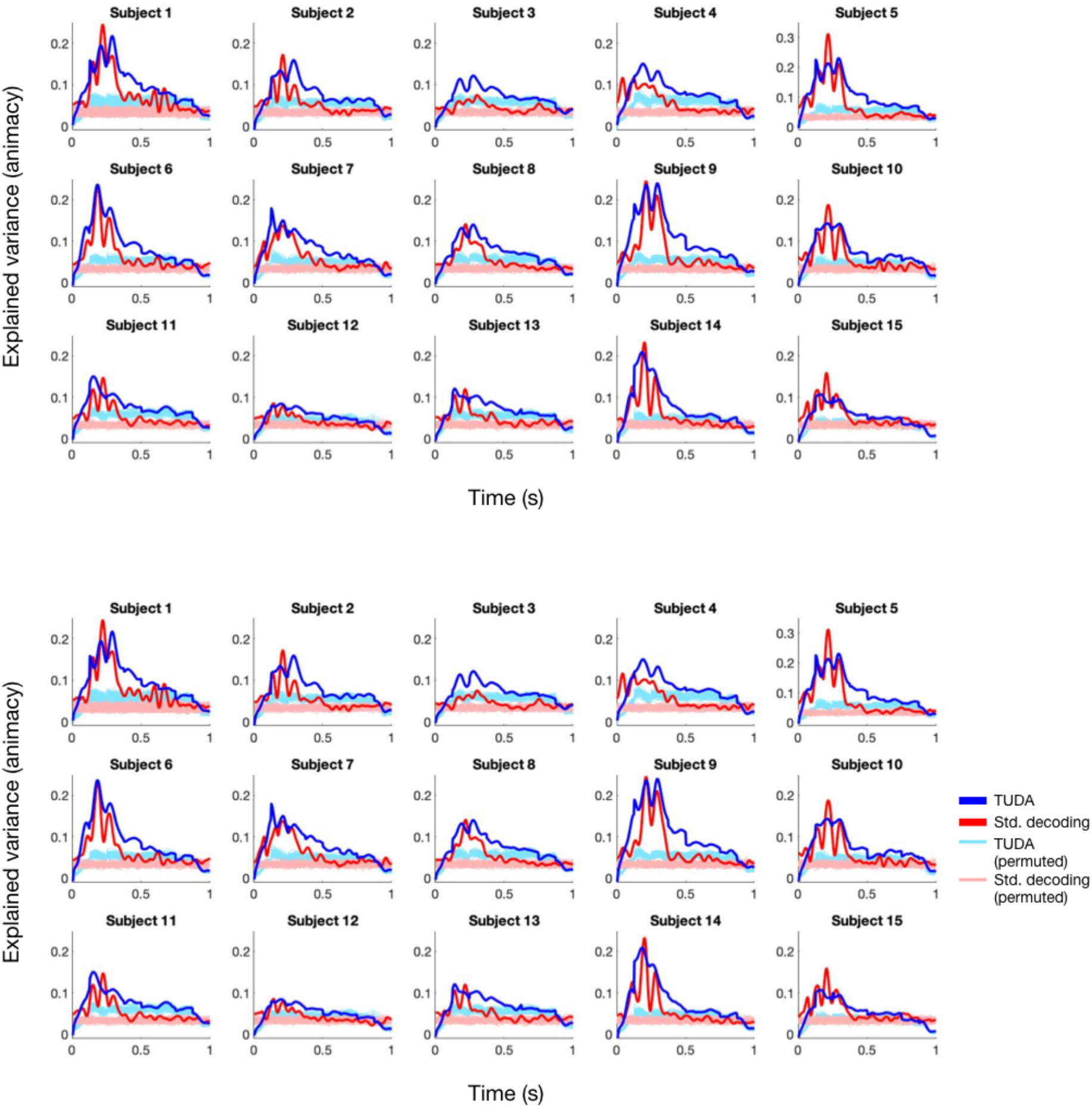
Subject by subject cross-validated decoding of size (top) and animacy (bottom).

